# Gut microbiome-associated adaptability of suckermouth catfish inhabiting the polluted Buriganga River: insights into heavy-metal-remediating microorganisms

**DOI:** 10.64898/2026.11.03.763882

**Authors:** Sujan Dey, Sumaita Alam Sikder, Mohsina Rahman Epa, Sujan Kanti Das, Md. Zobaidul Alam

**Author notes:** These authors contributed equally as first authors (Sujan Dey, Sumaita Alam Sikder).

## Abstract

The suckermouth catfish (SMCF), known for its pollutant tolerance and high invasiveness, is prevalent in the heavily contaminated Buriganga River in Bangladesh. This waterway, contaminated with a variety of pollutants, including heavy metals, has experienced the near-complete disappearance of native fish populations; however, the suckermouth catfish demonstrates remarkable adaptability in this environment. One objective of this study is to elucidate the role of the gut microbiome in SMCFs’ adaptation to this severely polluted habitat. We performed next-generation sequencing (NGS) of gut contents from SMCFs to analyze the gut microbial profiles of the fish samples. The results indicated that the suckermouth catfish’s gut is enriched with microbial communities, including *Paraclostridium*, *Klebsiella*, *Staphylococcus*, *Bacillus*, and *Pseudomonas*, which have the potential to remediate pollutants, including heavy metals. The subsequent aim of this research is to isolate and identify bacteria from the SMCF gut capable of remediating multiple heavy metals. We isolated three isolates with the potential to remediate multiple heavy metals and identified them as *Staphylococcus pasteuri* SD, *Morganella morganii* SAS1, and *Sporosarcina saromensis* SAS using partial 16S rRNA gene amplicon sequencing and conventional techniques. All isolates showed distinct capabilities for remediating multiple heavy metals. Notably, *S. pasteuri* showed the highest lead (Pb) bioaccumulation (36.28%), S. saromensis showed the highest chromium (Cr) uptake (39.22%), and *M. morganii* showed the highest nickel (Ni) uptake (35.90%). Furthermore, these isolates exhibited remarkable metal-tolerance profiles, with maximum tolerable concentrations ranging from 125 ppm to 1000 ppm, and sustained healthy growth under metal-supplemented conditions throughout a 48-hour incubation period. This validates their potential to remediate multiple heavy metals. Collectively, the results underscore the ecological adaptability of suckermouth catfish in polluted aquatic environments, which is associated with their gut microbiome, and highlight the bioremediation potential of these gut-derived microbes as effective agents for mitigating heavy metal contamination.

## Introduction

The suckermouth catfish (SMCF), classified within the order Siluriformes and the family Loricariidae, is a substantial and diverse group comprising approximately 90 genera and 846 documented species, originating from South and Central America [1]. Notably, the genera *Hypostomus* and *Pterygoplichthys* are frequently dominant in Asian regions. Because of their distinctive appearance and algae-cleaning capabilities, SMCF are widely kept as ornamental fish in aquaria; however, they are not generally consumed as food [2]. Reports indicate that introducing SMCF into new ecological environments can adversely affect ecosystem functioning, biodiversity, national economies, and public health [3]. These species pose a significant threat to native fish populations in the water bodies into which they are introduced, primarily because of their unique feeding behaviors, reproductive strategies, and ability to establish persistent populations [4,5]. In addition to their well-documented predation pressure, these catfish are also recognized for their durable external armor and remarkable capacity to withstand low levels of dissolved oxygen [6]. Their ability to thrive and proliferate in heavy-metal-contaminated environments has also been documented [7]. In the 1980s, these fish were imported into Bangladesh for trading purposes. Currently, SMCFs are rapidly proliferating in certain freshwater habitats, including rivers, because of their invasive nature, high reproductive rates, and ability to survive in polluted environments. Floodplains within the Ganges-Brahmaputra River drainage basin in northern and southwestern Bangladesh have been overrun by suckermouth catfish [8]. The aggressive behaviors, survival skills, and omnivorous diet of these fish raise concerns regarding the sustainability of local fisheries and the health of aquatic ecosystems. Their exceptional adaptability to highly polluted water bodies worldwide underscores the ecological threat they pose [9]. In response, the Government of Bangladesh mandated a prohibition on the cultivation, capture, and trade of suckermouth catfish in September 2022 [10].

The Buriganga River, one of Bangladesh’s principal waterways, is severely contaminated by a wide range of pollutants. Alongside industrial growth, a significant volume of household waste has been discharged into the river, markedly increasing pollution levels [11]. Over the past three decades, numerous textile and tannery industries have been established along the riverbank [12]. Research has documented that the Buriganga is polluted by various contaminants, including heavy metals, plastics, oil, dead animals, sewage, chemical effluents from manufacturing facilities, and waste from households and medical facilities. Each day, hundreds of tons of solid waste are generated in Dhaka city, most of which ultimately enters the river [13]. Heavy metals are non-biodegradable and can bioaccumulate through the food chain, posing serious risks to human health and the environment, even from distant sources of contamination [14]. Previous studies have reported multiple heavy metals, such as Cadmium (Cd), Chromium (Cr), Lead (Pb), and Arsenic (As), at most sampling sites along the Buriganga [15]. Some concentrations of these metals exceed recommended safety thresholds, making the fish, sediment, and river water unsafe for ecosystem integrity and public health [16]. The heavily contaminated Buriganga ecosystem presents a significant threat to aquatic flora and fauna, including edible fish species. Nevertheless, Suckermouth Catfish (SMCF) can survive and reproduce extensively in this polluted environment, whereas edible fish are present only in very limited numbers. The reasons for SMCF’s remarkable adaptability in such a hostile aquatic setting remain uncertain, especially as other edible fish species in the area face extinction risk. Furthermore, it is unclear whether the microbial communities in the SMCF gut contribute to its survival. One objective of this study is to characterize the gut microbial community profiles of Suckermouth Catfish collected from highly polluted sites along the Buriganga River.

Heavy metal contamination occurs when toxic metals such as chromium, lead, cadmium, nickel, mercury, zinc, and copper accumulate in the environment to hazardous levels, jeopardizing soil and aquatic ecosystems [17,18]. Heavy metals (HMs) influence soil microbiology by disrupting interactions between microorganisms and plants in the rhizosphere, thereby affecting soil characteristics, plant development, vegetation, and agricultural productivity. Exposure to heavy metals in aquatic ecosystems induces oxidative stress and interferes with normal protein folding and physiological functions in both vertebrates and invertebrates [19,20]. To remediate heavy metal pollution, strategies such as immobilization, extraction, sequestration, active mixing, or microbial bioremediation may be employed [21]. Among these, bioremediation has emerged as an economically viable method, offering an effective way to mitigate pollution and rehabilitate contaminated sites by degrading toxic and recalcitrant compounds, including heavy metals [22]. Bioremediation of soils contaminated with heavy metals using microorganisms or functional biocatalysts is a growing area within biotechnology. This process benefits from integrating advanced biological sciences, including genomics, transcriptomics, and proteomics, with synthetic biology [23]. Such interdisciplinary approaches facilitate the design and optimization of microbial systems that can efficiently mitigate heavy metal contamination [24]. Distinct microbial communities exhibit unique metabolic capabilities: some microorganisms interact with toxic metals to produce organic compounds, whereas others promote the formation of nanoparticles that reduce heavy metal toxicity [25]. Additionally, nanotechnology-based materials, due to their high surface area and reactivity, have been explored for micro-remediation of heavy metals in soils [26].

In summary, removing heavy metals (HM) using bioremediation techniques is a promising and rapidly advancing field of research, offering substantial potential to discover novel bacteria capable of comprehensive and efficient bioremediation. Consequently, another objective of this study is to isolate gut microbes from SMCF and evaluate their potential for heavy-metal bioremediation. Overall, this study was designed with two specific aims: (i) to elucidate the role of the gut microbial profile of SMCF in their remarkable adaptability to a heavily polluted environment and (ii) to isolate and characterize potential multiple heavy metal-tolerant (HMT) bacterial strains from the gut of SMCF with enhanced capabilities for environmental heavy metal remediation.

## Materials and Methods

### Water and fish sample collection and sample preparation

Four sampling stations along the Buriganga River were identified: Station1 at Postogola Bridge (23.6869° N, 90.426847° E), Station 2 at Fatullah-Pagla (23.6387° N, 90.4836° E), Station 3 at Kamrangirchar (23.7153° N, 90.3736° E), and Station 4 at Sadarghat (23.725° N, 90.405° E). Fish and water samples were collected at each site. Water samples were transported immediately to the laboratory for physicochemical analysis. Concurrently, a total of 12 suckermouth fish samples (SMCF), later identified as *Pterygoplichthys disjunctivus*, were collected at four sampling stations (S1 Fig.). Subsequently, the fish samples were weighed and measured for length. Then, the fish were dissected under sterile conditions to obtain gut, muscle, and liver tissues. The gut was carefully incised with sterile scissors and forceps into small pieces, then placed in sterile Falcon tubes for further analysis.

### DNA extraction and NGS analysis of fish gut content

Genomic DNA was extracted from fish gut contents using the DNeasy PowerSoil Pro Kit (Cat. no. 47014) following the manufacturer’s protocol. The extracted DNA was quantified using a NanoDrop spectrophotometer (Thermo Fisher Scientific, USA), and DNA integrity was assessed by electrophoresis on a 1% agarose gel. For microbial community analysis, PCR amplification of the extracted DNA was performed targeting the V3-V4 region of the 16S rRNA gene using the Universal linker primers 341F (5′-CCTAYGGGRBGCASCAG-3′) and 806R (5′-GGACTACHVGGGTWTCTAAT-3′), which carry distinct sample barcodes [27]. PCR amplification was conducted in accordance with methodologies previously described by other researchers. In summary, all PCR reactions were performed in a 30 mL volume buffer containing Promega G2 Green Master Mix, forward and reverse primers, template DNA, and nuclease-free water. The amplification protocol consisted of an initial denaturation at 98°C for 1 minute, followed by 30 cycles of denaturation at 98°C for 10 seconds, annealing at 50°C for 30 seconds, and extension at 72°C for 30 seconds, culminating in a final extension at 72°C for 5 minutes. PCR products were purified using the GeneJET Gel Extraction Kit, quantified with a Qubit fluorometer according to the manufacturer’s instructions, and validated by analysis on a 2% agarose gel. The sequencing library was assembled using equimolar amounts of purified amplicons, following the Illumina TruSeq library preparation protocol. The final libraries were sequenced on the Illumina NovaSeq 6000 platform in paired-end (2 × 250 bp) mode to generate high-throughput reads for downstream bioinformatics analysis. Chimeric sequences were removed with VSEARCH, and sequences with ≥97% identity were clustered into operational taxonomic units (OTUs) using QIIME 2. Taxonomic classification was performed using the SILVA 138 reference database [27–30].

### Physicochemical parameters analysis of water samples

At each sampling location, temperature and pH were measured with a handheld laboratory thermometer (Mercury 305 MM, Zeal, England) and a calibrated pH meter (HI98107, Hanna Instruments, Germany), respectively. Electrical conductivity (EC) was measured with an EC meter (EC-214, Hanna Instruments Ltd., UK). Other water quality parameters, including biological oxygen demand (BOD), chemical oxygen demand (COD), dissolved oxygen (DO), total dissolved solids (TDS), and total suspended solids (TSS), were analyzed following standardized protocols [31–33]. Anion concentrations were determined by ion chromatography. All experiments were performed in triplicate. To assess significance, a one-way ANOVA with three replicates was used at a threshold of p < 0.05.

### Water quality index (WQI)

To evaluate overall water quality, the Water Quality Index (WQI) was calculated using the methodology outlined in a previous study [34,35]. Nine parameters—pH, BOD, COD, DO, TDS, TSS, nitrate, chloride, and sulfate—were incorporated to determine the WQI. The calculation of the Water Quality Index (WQI) used the following formula:

WQI = ∑QiWi

where Qi (water quality rating) = 100 × (Va-Vi)/ (Vs-Vi); Va= actual value present in the water sample; Vi= ideal value (0 for all parameters except pH and DO, which are 7.0 and 14.6 mg/1, respectively) and Vs= Standard value.

Wi (Unit weight) = K/Sn

where K (constant) =1/ (1/Vs1+1/Vs2+1/Vs3+1/Vs4+………+1/Vsn) and Sn= ‘n’ number of standard values.

### Quantitative analysis of heavy metals in river water and fish muscle

To perform heavy metal analysis of the fish and water samples, the collected water and fish muscle samples were acid-digested with nitric acid (HNO₃), then filtered through Whatman No. 42 filter paper. The final volume was adjusted to 100 mL with distilled water. The analysis was conducted using an atomic absorption spectrophotometer (ICE 3000 AA system, Thermo Fisher Scientific) with an air-acetylene flame. To determine heavy metal concentrations in the water samples and fish muscle, each measurement was performed in triplicate, and the resulting values were statistically analyzed by calculating the mean and standard deviation to ensure accuracy and reproducibility. The wavelength of maximum absorption, or λmax, was used for all spectroscopic measurements of the sample solutions and standard metal solutions [36–38].

### Screening of multiple heavy metal-remediating bacterial isolates

Suckermouth catfish gut samples were serially diluted (10⁻¹ to 10⁻⁸). LB agar was prepared by supplementing it with one of the following heavy metals: CdCl₂·H₂O, NiSO₄·6H₂O, Pb(C₂H₃O₂)₂·3H₂O, CuSO₄·5H₂O, FeSO₄·7H₂O, or K₂Cr₂O₇ to a final concentration of 100 ppm. 100 μL from each dilution was spread onto heavy metal-supplemented LB agar plates and incubated at 30°C for 24 hours. After incubation, colony formation indicated metal tolerance. Each isolate was then tested for tolerance to multiple heavy metals (HMs) by streaking onto each metal-supplemented LB agar plate and incubating at 30°C for 24 hours [39].

### Maximum tolerable concentration (MTC) of heavy metals by the isolates

The Maximum Tolerable Concentration (MTC) of the isolates was determined to assess their potential for heavy-metal remediation. In this study, MTC is defined as the highest metal concentration that allows visible growth after incubation. One set of Mueller-Hinton broth was prepared by supplementing it with the salts of the respective heavy metals (CdCl₂·H₂O, NiSO₄·6H₂O, Pb(C₂H₃O₂)₂·3H₂O, CuSO₄·5H₂O, FeSO₄·7H₂O, or K₂Cr₂O₇) to a final concentration of 16,000 ppm. Another Mueller-Hinton broth was prepared without heavy metal supplementation. The MTC assay was conducted in a sterile microtiter plate. In the first well of the plate, 100 µL of metal-supplemented Mueller-Hinton broth was added, and the 2nd to 10th wells were dispensed with 50 µL of metal-free Mueller-Hinton broth. Thereafter, serial dilutions of the metal-supplemented Mueller-Hinton broth were prepared by successive transfers of 50 µL from the 1st well to the 10th, thereby creating a gradual decrease in concentration across the wells. Finally, 50 µL of broth was discarded from the 10th well. 10 µL of bacterial cultures (OD₆₀₀ = 0.08–0.1) was inoculated into each well. The 11^th^ well of the plate was maintained as a negative control with 50 µL of metal-supplemented Mueller-Hinton broth without inoculation. The 12^th^ well served as a positive control and contained 50 µL of metal-free Mueller-Hinton broth inoculated with bacteria. Plates were sealed and incubated at 30°C for 48 hours. After incubation, 20 µL of 0.1% TTC (2,3,5-Triphenyltetrazolium chloride) was added to each well, followed by a 2-hour incubation at 30°C. Red coloration indicated metabolic activity. Growth responses were categorized as: no growth (high toxicity), reduced growth (moderate toxicity), or normal growth (low/no toxicity) by observing visual color change [40,41].

### Determination of growth curves of bacterial isolates

To assess the effect of heavy metals on bacterial growth, a growth curve was obtained. By comparing the growth patterns under heavy metal exposure and control conditions, the influence of heavy metals on the different phases of bacterial growth was determined. For this purpose, LB broth was prepared by supplementing it with the salts of the respective heavy metals: CdCl₂·H₂O, NiSO₄·6H₂O, Pb(C₂H₃O₂)₂·3H₂O, CuSO₄·5H₂O, FeSO₄·7H₂O, or K₂Cr₂O₇, each at a final concentration of 100 ppm. 150 ml of metal-supplemented LB broth was dispensed into a series of 250 ml conical flasks, inoculated with 100 µL of overnight-grown bacterial culture, sealed properly, and incubated at 30 °C for 48 hours under shaking conditions. The control flask contained 150 ml of metal-free LB broth inoculated with 100 µL of inoculum. Growth was monitored by measuring OD₆₀₀ at 4-hour intervals during the incubation period up to 48 h. OD vs. time plots were generated for each condition [39].

### Bioremediation assay

To conduct the bioremediation assay, LB broth was prepared by supplementing it with the salts of the respective heavy metals: CdCl₂·H₂O, NiSO₄·6H₂O, Pb(C₂H₃O₂)₂·3H₂O, CuSO₄·5H₂O, FeSO₄·7H₂O, or K₂Cr₂O₇, each at a final concentration of 100 ppm. 150 mL of metal-supplemented LB broth was dispensed into a series of 250 mL conical flasks, inoculated with 150 µL of overnight-grown bacterial culture, sealed properly, and incubated at 30 °C for 48 hours under shaking conditions. The initial and final concentrations of the metal in the culture medium were measured by Atomic Absorption Spectroscopy (AAS) before and after incubation. For this, 50 ml of culture from each set of samples was centrifuged, the supernatants were collected, and the samples were analyzed by Atomic Absorption Spectroscopy (AAS). The percentage of metal bioremediation or accumulation by each isolate was calculated as follows[39,40].

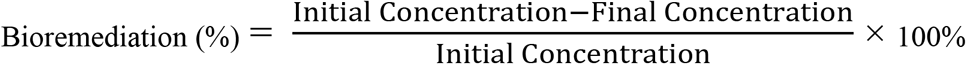

Here, the initial HM concentration is the heavy metal concentration in the culture medium before incubation, and the final HM concentration is the heavy metal concentration in the culture medium after incubation.

### Identification of heavy metal-remediating microbial isolates

Following primary screening, three prospective microbial isolates, designated B1, B2, and B3, capable of remediating multiple heavy metals (HMs), were identified through partial sequencing of 16S rRNA gene amplicons. The 16S rRNA gene amplicons of these isolates were amplified to nearly full length using universal primers 27F (5’-AGAGTTTGATCNTGGCTCAG-3’) and 1492R (5’-GCTTACCTTGTTACGACTT-3’), targeting approximately 1400 base pairs of the gene [42]. A total of 20-μL PCR reaction volume contained 10 μL GoTaq G2 Hot Start Master Mix (Promega), 1 μL template DNA, 1 μL of each primer (10 to 20 pM), and 7μL of PCR-grade water. PCR amplification procedures commenced with an initial denaturation at 95°C for 3 minutes, followed by 35 cycles of denaturation at 95°C for 30 seconds, annealing at 48°C for 30 seconds, extension at 72°C for 90 seconds followed by final extension at 72°C for 5 minutes and cooling at 4°C. Prior to sequencing the PCR products were purified using Wizard SV Gel and PCR Clean-Up System (Promega). Sequencing of the purified PCR amplicons was performed utilizing the BigDye™ Terminator v3.1 kit (Applied Biosystems, Thermo Fisher Scientific K.K.) and analyzed by the 3730xl DNA Analyzer in accordance with the manufacturer’s protocol. Obtained sequences were further processed using BioEdit software and deposited in the National Center for Biotechnology Information (NCBI) Data Bank with accession numbers PZ182827.1, PZ179984.1 and PZ492667.1. Furthermore, the three isolates were characterized based on their morphological, physiological, cultural, and biochemical attributes (Table S3).

### Phylogenetic analysis

Using BLAST (megablast) through the NCBI GenBank database, sequence similarity analysis of 16S rRNA was conducted for each isolate. The top-scoring hit was used to assign genus and/or species identity. The sequences of 16S rRNA were aligned using the multiple sequence alignment program CLUSTAL-W. Phylogenetic and molecular evolutionary analyses were processed through the molecular evolutionary genetics analysis software MEGA 12 [43,44].

### Statistical analysis

All experiments were conducted with at least three replicates. To assess statistical significance, we performed a one-way ANOVA with three replicates using IBM SPSS v. 26.

## Results

### Current pollution status of Buriganga River and water quality index (WQI)

To assess the current pollution levels in the Buriganga River and to determine its suitability as a sampling site for the collection of SMCF, the Water Quality Index (WQI) (Table 1) was computed based on the measured values of various physicochemical parameters of the river water, as detailed in S1 and S2 Tables. The physicochemical analysis revealed numerous indicators of severe pollution, as all measured parameters exceeded WHO standards at most stations. Additionally, the WQI in Table 1 provides a comprehensive overview of the Buriganga River’s current pollution status by synthesizing a range of physicochemical indicators into a single score. The WQI values obtained at the four sampling stations indicate statistically significant (P < 0. 0.05) pollution across all locations. Station 2 had the highest WQI (203.51), categorizing the water quality as extremely polluted and indicating substantial deterioration. Stations 1 and 3 showed WQI values of 153.25 and 146.44, respectively, both falling within the very poor to extremely polluted categories. Station 4, with a WQI of 120.91, also reflects substandard quality, though conditions are marginally less severe than at the other sites. Heavy metal concentrations in the Buriganga River water were likewise analyzed to assess the extent of metal pollution and evaluate the biological implications for aquatic life. Eight heavy metals-Cobalt (Co), Cadmium (Cd), Chromium (Cr), Lead (Pb), Manganese (Mn), Copper (Cu), Zinc (Zn), and Nickel (Ni)-were examined. Table 2 shows that heavy metal concentrations at the four sampling sites exceeded the WHO permissible limits for Cd, Cr, Pb, and Ni. Although the WHO has no specific standard for Co, its elevated concentration at station 4 (0.33 ± 0.05 ppm) raises ecological concerns. Conversely, Mn, Cu, and Zn concentrations remain within acceptable ranges. These elevated WQI values, coupled with increased heavy metal concentrations in the river water, collectively demonstrate the severe pollution of the Buriganga River and thereby confirm its suitability for collecting SMCF for this study.

**Table 1.** Water Quality Index of the River Water Samples.

|  | **Water Quality Rating (qi)** |  |  |  | **Unit Weight (wi)** | **Water Quality Index (wiqi)** |  |  |  |
|---|---|---|---|---|---|---|---|---|---|
| **Parameter** | **Station 1** | **Station 2** | **Station 3** | **Station 4** |  | **Station 1** | **Station 2** | **Station 3** | **Station 4** |
| **pH** | 200 | 193.33 | 100 | 46.67 | 0.367 | 73.22 | 70.78 | 36.61 | 17.09 |
| **BOD** | 177.67 | 282 | 262 | 225.00 | 0.318 | 56.57 | 89.79 | 83.42 | 71.64 |
| **COD** | 64.00 | 169.60 | 90 | 129.88 | 0.159 | 10.19 | 27.00 | 14.33 | 20.68 |
| **TDS** | 139.23 | 129.11 | 77.25 | 111.00 | 0.003 | 0.45 | 0.41 | 0.25 | 0.36 |
| **TSS** | 534 | 584 | 600.50 | 650.3 | 0.003 | 1.71 | 1.87 | 1.92 | 2.08 |
| **EC** | 71.4 | 71.33 | 66.07 | 60.6 | 0.002 | 0.11 | 0.11 | 0.11 | 0.10 |
| **Nitrate** | 17.66 | 10.26 | 8.262 | 16.15 | 0.062 | 1.10 | 0.64 | 0.51 | 1.01 |
| **Chloride** | 11.23 | 9.86 | 10.65 | 12.396 | 0.012 | 0.14 | 0.12 | 0.13 | 0.15 |
| **Sulfate** | 31.41 | 26.90 | 16.94 | 33.385 | 0.012 | 0.40 | 0.34 | 0.21 | 0.42 |
| **<sup>a</sup>WQI** |  |  |  |  |  | 153.25 | 203.51 | 146.44 | 120.90 |

<sup>a</sup>WQI>100; poor water quality [45]

**Table 2.** Heavy metal concentration of the river water samples (ppm)

| **Metal** | **Station 1** | **Station 2** | **Station 3** | **Station 4** | **Acceptable limit(WHO)[46]** |
|---|---|---|---|---|---|
| **Co** | 0.18±0.10 | 0.16±0.06 | 0.20±0.10 | 0.33±0.05 | - |
| **Cd** | 0.08±0.01 | 0.05±0.02 | 0.09±0.01 | 0.07±0.02 | 0.003 |
| **Cr** | 0.09±0.07 | 0.09±0.05 | 0.12±0.02 | 0.14±0.05 | 0.05 |
| **Pb** | 0.11±0.01 | 0.15±0.06 | 0.21±0.10 | 0.14±0.04 | 0.01 |
| **Mn** | 0.07±0.01 | 0.13±0.06 | 0.09±0.01 | 0.31±0.03 | 0.4 |
| **Cu** | 0.12±0.01 | 0.22±0.02 | 0.25±0.04 | 0.37±0.06 | 2.00 |
| **Zn** | 0.31±0.12 | 1.39±0.82 | 0.45±0.03 | 0.51±0.03 | 3.00 |
| **Ni** | 0.13±0.01 | 0.16±0.01 | 0.21±0.02 | 0.48±0.04 | 0.07 |

### Microbial community profile of the gut content of the SMCFs

The microbial community profile in the gut of SMCFs was determined using next-generation sequencing (NGS) of 16S rRNA gene amplicons, as shown in Fig 1. In SMCFs, the phylum Firmicutes was most prevalent, comprising 58%-70% of the microbial community, followed by 25%-35% Proteobacteria and 2%-6% Actinobacteriota. Within Firmicutes, *Paraclostridium* (order Peptostreptococcales) was the most abundant, with a relative abundance of 29.7% to 39.5%. In addition to *Paraclostridium*, other Firmicutes genera present in the SMCF included *Bacillus* (15.4% to 26.7%), *Staphylococcus* (3.5% to 9.8%), and *Sporosarcina (*1.6% to 3.4%), classified within the order Bacillales, and *Enterococcus* (1.1% to 2.5%), within the order Lactobacillales.

**Fig 1.**
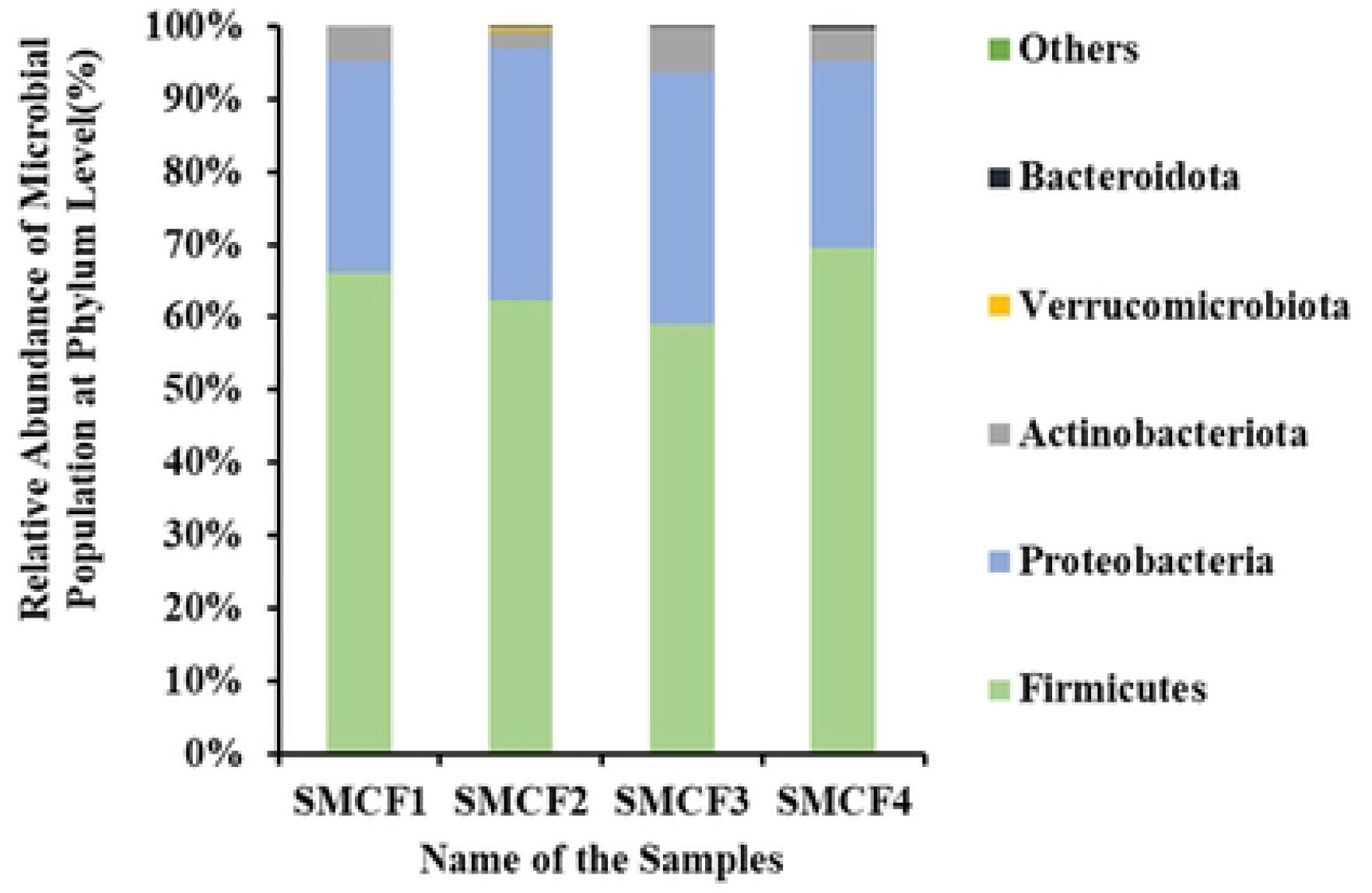
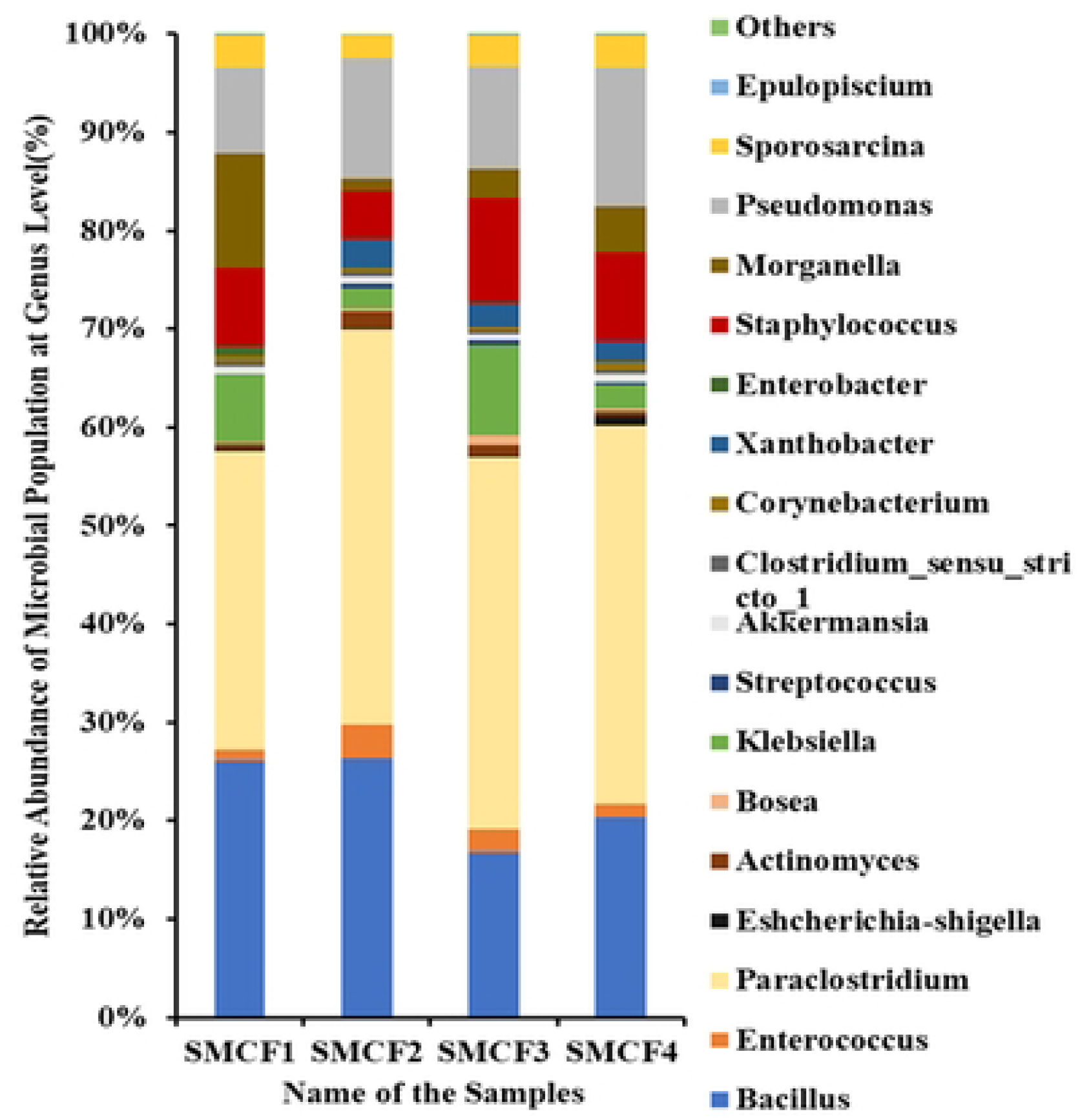
Relative abundance of the microbial profile at the phylum (A) and genus (B) levels in suckermouth catfish (SMCF). SMCF1, SMCF2, SMCF3, and SMCF4 represent suckermouth catfish collected from stations 1, 2, 3, and 4, respectively. The “Others” category comprises a diverse set of additional genera detected only at negligible abundances.

The microbial community in the SMCF also included representatives of the phylum Proteobacteria, genera such as *Pseudomonas* (8.9%-14.5%), *Morganella* (2.6%-11.9%), and *Klebsiella* (1.5%-8.4%) all belonging to the class Gammaproteobacteria, which were identified as comprising a substantial proportion of this phylum. Within the phylum Actinobacteriota, Actinomyces and Corynebacterium accounted for 0.4%-1.2% and 0.3%-0.9%, respectively. Finally, the “others” category encompassed a diverse array of additional genera identified only at negligible abundances. This suggests that although individual genera in this group contributed minimally to the overall microbial composition, their collective presence may be important for SMCF adaptability in polluted environments. Bacteria in SMCF show promising prospects for heavy-metal bioremediation. Specifically, SMCF harbors higher abundances of contaminant-remediating bacterial strains, including *Paraclostridium, Klebsiella, Staphylococcus, Bacillus*, and *Pseudomonas*, which can bioremediate heavy metals and other pollutants.

Shannon diversity index for the SMCF samples is 1.74±0.16, 1.78±0.04, 1.83±0.13, and 1.88±0.07, with an evenness of 0.66, 0.60, 0.66, and 0.64 for SMCF 1, SMCF 2, SMCF 3, and SMCF 4, respectively. Gut microbial diversity index did not differ significantly (p > 0.05) among fish samples collected from four locations along the Buriganga River. These results indicate that SMCF gut microbial community structure across the Buriganga River is relatively stable.

### Heavy metals in SMCF muscle

We measured heavy metal concentrations in SMCF to assess the fish’s capacity for heavy metal bioaccumulation. Table 3 shows heavy metal concentrations in suckermouth catfish samples SMCF1, SMCF2, SMCF3, and SMCF4, collected from four locations along the Buriganga River. Lead (Pb) levels are notably elevated across all samples, with SMCF4 (2.03 ± 0.06 ppm) exhibiting the highest concentration, which significantly exceeds the World Health Organization (WHO) guideline of 0.3 ppm. Cadmium (Cd) concentrations are particularly alarming in SMCF2 (0.39 ± 0.27 ppm), SMCF3 (0.44 ± 0.01 ppm), and SMCF4 (0.77 ± 0.07 ppm), all surpassing the permissible limit of 0.05 ppm, indicating considerable industrial influence. Manganese (Mn) also exceeds the safe threshold of 1.0 ppm in all samples, notably in SMCF2 (2.74 ± 0.26 ppm) and SMCF4 (2.54 ± 0.1 ppm), reflecting possible inputs from industrial effluents. Nickel (Ni) concentrations exceed the WHO permissible limit of 0.5 ppm in SMCF1 (1.07 ± 0.39 ppm) and SMCF3 (.87 ± 0.03 ppm). Although copper (Cu) and zinc (Zn) levels remain within acceptable limits, copper in SMCF3 (28.17 ± 0.2 ppm) approaches the upper boundary of 30.0 ppm, while zinc concentrations are highest in SMCF1 (15.89 ± 0.38 ppm), yet remain well below the limit of 100.0 ppm. Chromium (Cr) concentrations are within safe limits across all samples, with SMCF2 (0.98 ± 0.07 ppm) close to the 1.0 ppm threshold. Cobalt (Co) levels are relatively low and remain within the World Health Organization guideline of 0.5 ppm.

**Table 3:** Heavy metal concentrations in fish muscles (mg\kg)

| **Sample** | **SMCF1** | **SMCF2** | **SMCF3** | **SMCF4** | **Acceptable Limit (FAO)[47]** |
|---|---|---|---|---|---|
| **Co** | 0.043± 0.02 | 0.28 ± 0.16 | 0.36±0.03 | 0.05±0.04 | 0.50 |
| **Cd** | 0.05 ± 0.04 | 0.39 ± 0.27 | 0.44±0.01 | 0.77±0.07 | 0.05 |
| **Cr** | 0.49 ± 0.37 | 0.98 ± 0.07 | 0.67±0.02 | 0.27±0.04 | 1.00 |
| **Pb** | 1.24 ± 0.17 | 1.77 ± 0.22 | 1.37±0.02 | 2.03±0.06 | 0.30 |
| **Mn** | 1.87 ± 0.07 | 2.74 ± 0.26 | 1.99±0.02 | 2.54±0.1 | 1.00 |
| **Cu** | 26.09 ±0.45 | 25.89 ±0.32 | 28.17±0.2 | 27.83±0.28 | 30.0 |
| **Zn** | 15.89± 0.38 | 13.45± 0.16 | 9.53±0.38 | 11.37±0.4 | 100.0 |
| **Ni** | 1.07 ± 0.39 | 0.4 ± 0.20 | 0.87±0.03 | 0.79±0.04 | 0.50 |

### Isolation of multiple heavy metal-remediating bacteria

After primary screening, we obtained 20 distinct isolates from the fish gut sample. Five of these isolates demonstrated pronounced tolerance to multiple heavy metals. Of these five, we selected isolates B1, B2, and B3 for further investigation based on their colony morphology, while excluding the remaining two from the study.

### Maximum tolerable concentration (MTC) of heavy metals by the isolates

The maximum tolerance concentrations (MTCs) of six heavy metals (Ni, Cu, Cr, Cd, Fe, and Pb) were assessed for each isolate to determine the highest concentrations at which growth could be sustained following a 48-hour incubation period. Fig 2 presents the maximum concentrations of these metals at which individual isolates exhibited growth. Isolate B1 had an MTC of 1000 ppm for Pb and 500 ppm for Ni, Cu, and Cr. Under Cd and Fe conditions, B1 had comparatively lower MTCs of 250 and125 ppm, respectively. Isolate B2 had MTCs of 1000 ppm for Cr and 500 ppm for Ni and Pb. Under Cu, Cd, and Fe conditions, B2 had comparatively lower MTCs ranging from 125 to 250 ppm. For isolate B3, the MTC for Ni and Cu was 1000 ppm, the highest value recorded for this isolate. This isolate had MTCs of 500 ppm for Cr, Cd, and Fe, whereas the MTC for Pb was 250 ppm.

**Fig 2.**
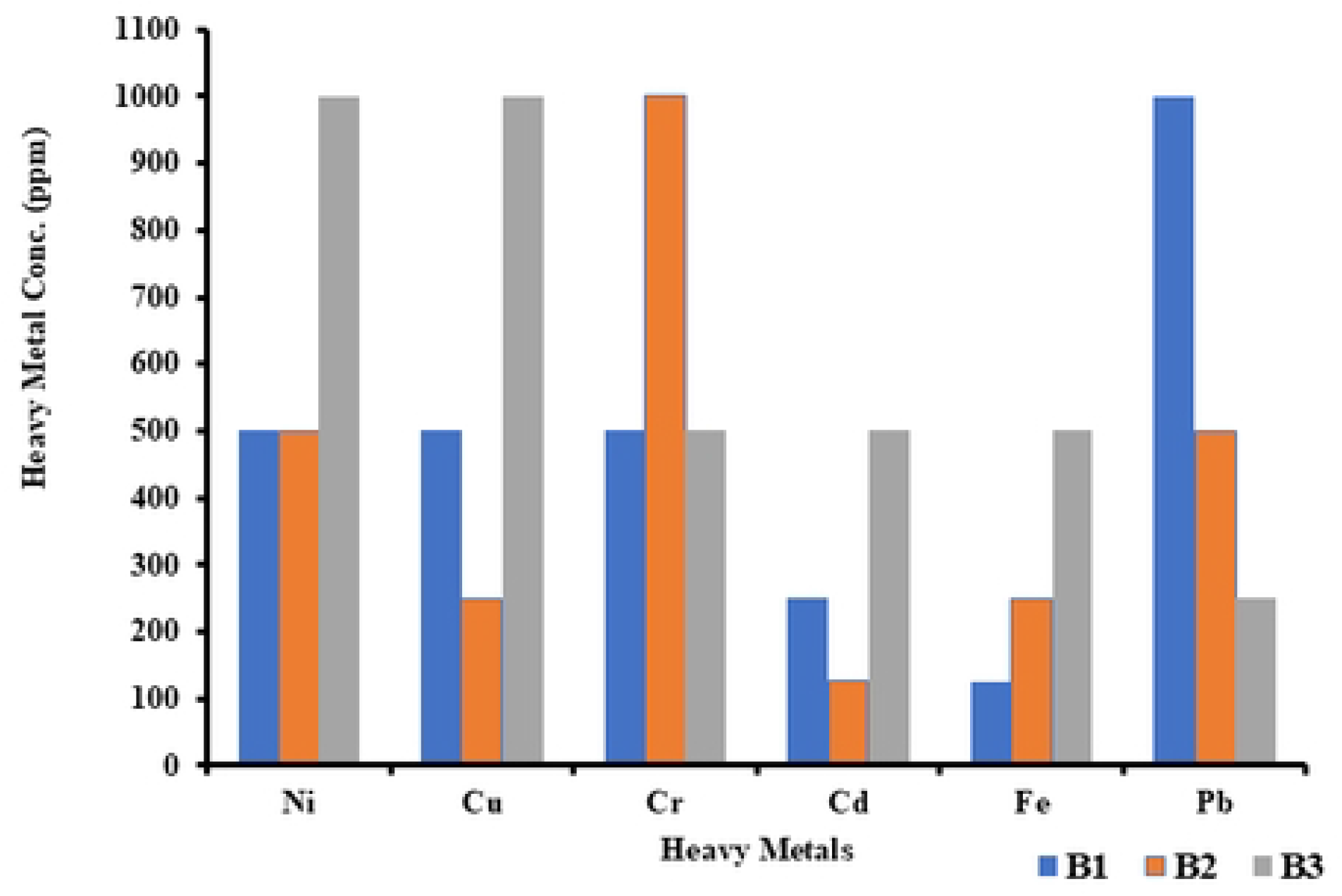
Maximum tolerable concentration of heavy metal salts in bacterial isolates: B1, B2, and B3. Heavy metal accumulation (Ni, Cu, Cr, Cd, Fe, Pb) was measured in three bacterial strains, B1 (Blue), B2 (Orange), and B3 (Grey). Bars represent mean heavy-metal concentrations (PPM), with error bars indicating variability across replicates. The x-axis shows the heavy metals tested in the experiment, while the y-axis represents their concentrations.

### Growth curve analysis of isolates

A growth curve was constructed to assess how heavy metals affect bacterial proliferation. Comparing growth patterns in the presence of heavy metals with those under control conditions revealed their effects on distinct bacterial growth phases. The growth curve in Fig 3 indicates that isolate B1 proliferated robustly under Ni, Cu, and Pb treatments, with high absorbance peaks, suggesting strong tolerance to heavy metals. Notably, this isolate showed the highest growth peak under Pb among all studied metals. Moderate growth was observed in the presence of Cr, Cd, and Fe. Overall, the isolate exhibited a distinct, differential growth pattern across heavy metal treatments. Likewise, the growth curve of isolate B2 (Fig 4) revealed enhanced tolerance to Pb, as the OD value remained elevated over a long incubation period, whereas under Ni and Cu conditions, it showed measurable growth but did not surpass the control level. However, the isolates showed (Fig 5) moderate growth under Cr-, Cd-, and Fe-supplemented conditions, which was lower than under the other heavy-metal treatments. From the growth curve shown in Fig 5, isolate B3 showed nearly the same level of growth under Ni-supplemented conditions as under the control condition. However, under other metal-supplemented conditions, growth was lower than growth under the control condition.

**Fig 3.**
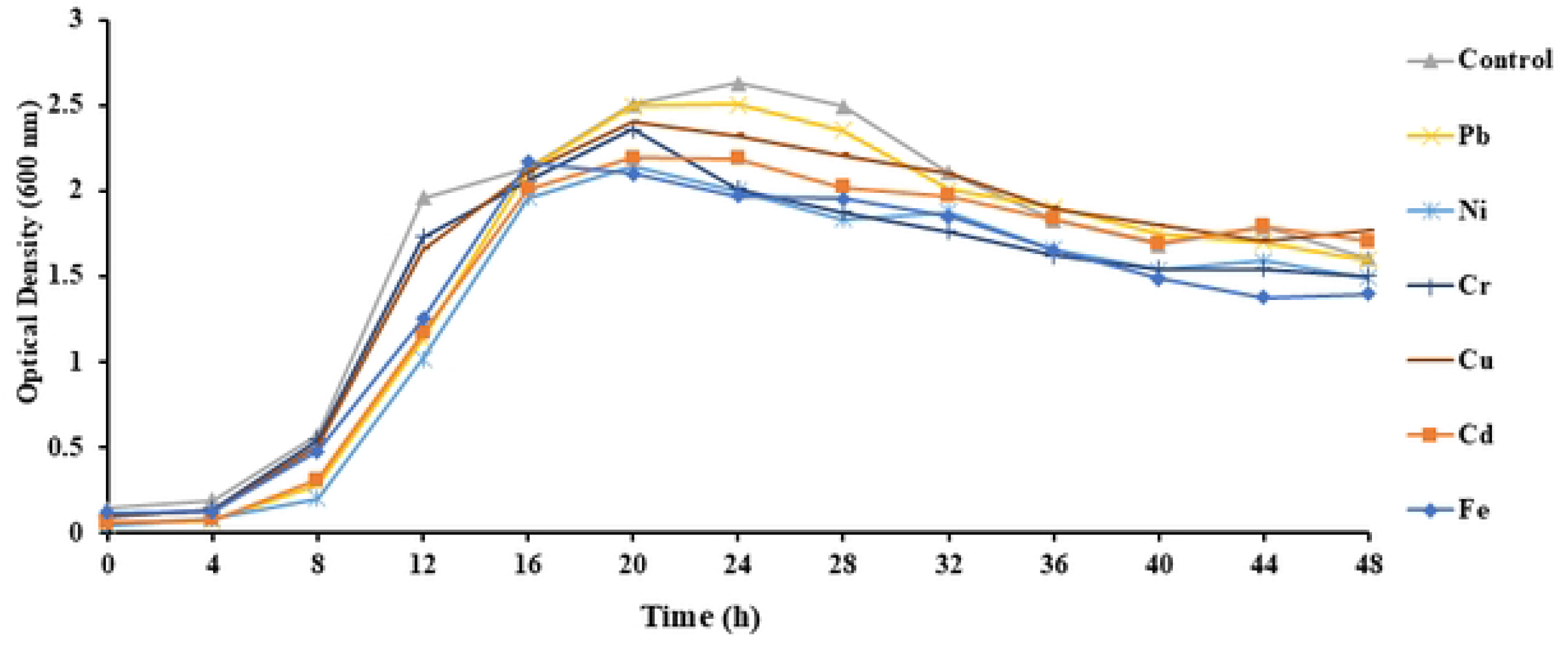
Growth curve of isolate B1. The x-axis represents time in 4-hour intervals over 48 hours, while the y-axis indicates OD absorbance values. Growth curves vary distinctly among treatments, reflecting differences in microbial responses to individual heavy metals. Treatments included LB broth supplemented with salts of different heavy metals (Ni, Pb, Cr, Cu, Cd, and Fe), and a control received only LB broth without heavy metal salt supplementation.

**Fig 4.**
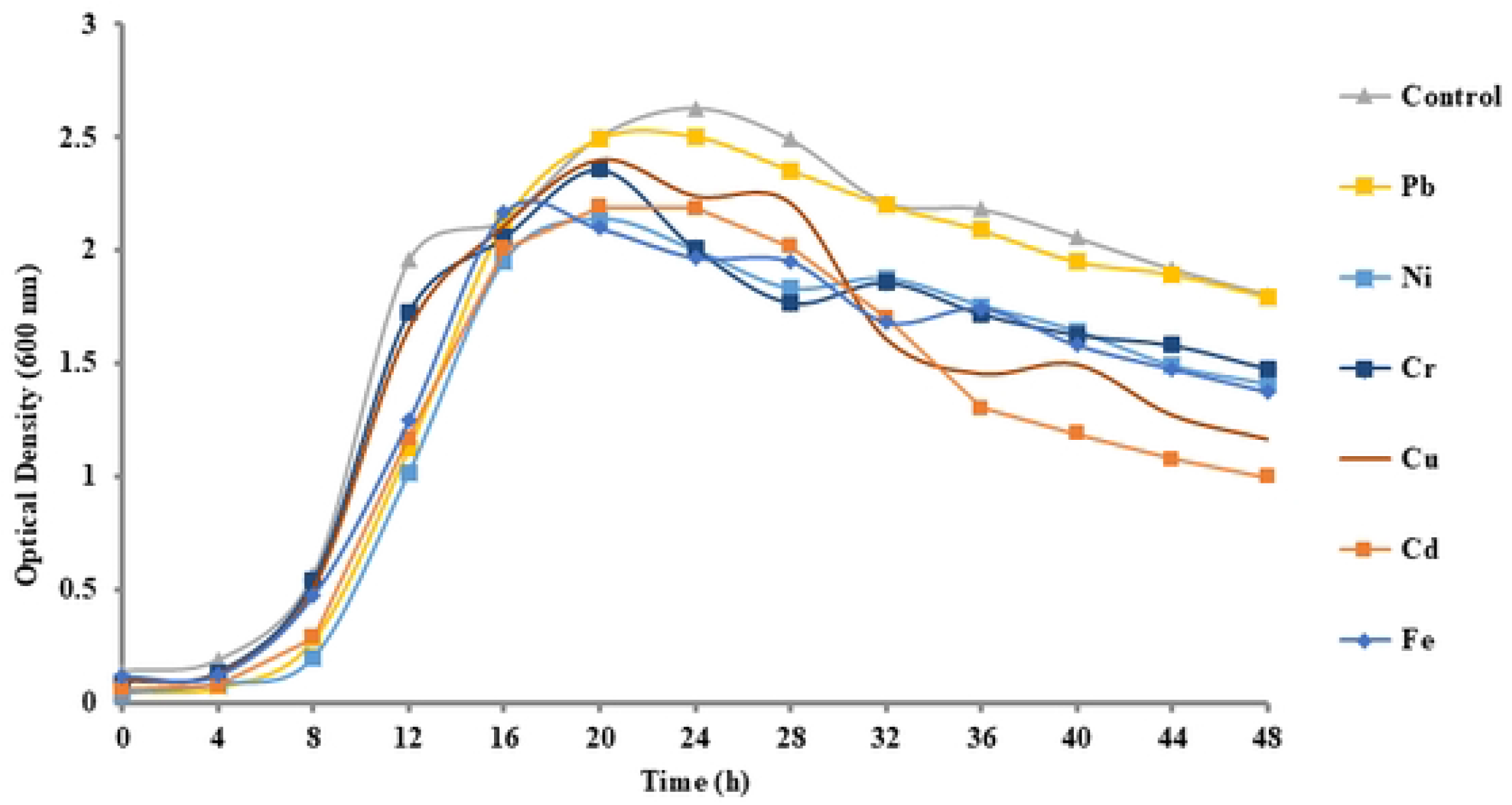
Growth curve of isolate B2. The x-axis represents time in 4-hour intervals over 48 hours, while the y-axis indicates OD absorbance values. Growth curves show distinct variations among treatments, reflecting differences in microbial response to individual heavy metals. Treatments included LB broth supplemented with salts of different heavy metals (Ni, Pb, Cr, Cu, Cd, and Fe), and a control received only LB broth without heavy metal salt supplementation.

**Fig 5.**
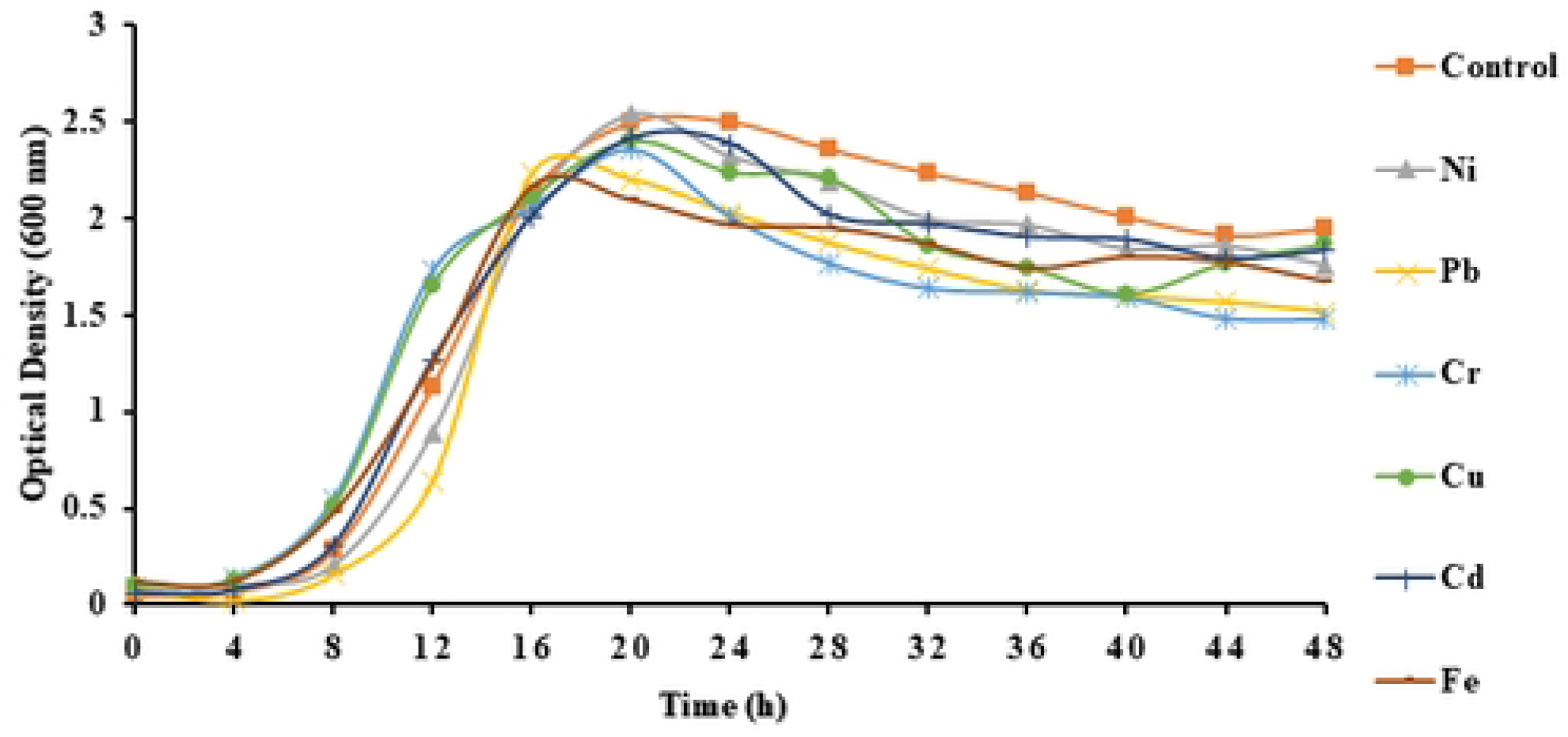
Growth curve of isolate B3. The x-axis represents time in 4-hour intervals over 48 hours, while the y-axis indicates OD absorbance values. Growth curves show distinct variations among treatments, reflecting differences in microbial response to individual heavy metals. Treatments included LB broth supplemented with salts of different heavy metals (Ni, Pb, Cr, Cu, Cd, and Fe), and a control received only LB broth without heavy metal salt supplementation.

### Bioremediation assay

After determining the MTCs of the isolates for heavy metals and analyzing their growth patterns in metal-supplemented LB medium, we evaluated the bacteria’s bioaccumulation capacity for various metals. Isolate B1 exhibited variable bioaccumulation efficiencies for the metals tested (Table 4). The highest uptake was observed for lead (Pb) at 36.28%, indicating strong tolerance to this metal. For copper (Cu) at 29.35% and nickel (Ni) at 27.48%, the isolate also displayed notable accumulation potential. Conversely, chromium (Cr) at 25.47% and cadmium (Cd) at 19.84% showed lower uptake, with the minimum uptake (16.30%) observed for iron (Fe). Isolate B2 (Table 5) demonstrated the highest bioaccumulation for chromium (Cr) at 39.22%, followed by nickel (Ni) at 33.18%. The bioaccumulation percentages for other metals in this isolate were as follows: lead (Pb) at 32.58%, iron (Fe) at 21.05%, copper (Cu) at 18.98%, and cadmium (Cd) at 18.57%. Lastly, isolate B3 (Table 6) exhibited the highest bioaccumulation for nickel (Ni) at 35.90%, followed by copper (Cu) at 30.65%. For the remaining metals, bioaccumulation levels were below 30%: chromium (28.40%), cadmium (27.59%), and iron (25.60%). Notably, this isolate showed the lowest lead (Pb) accumulation (21.30%).

**Table 4.** Bioaccumulation by isolate B1 (*Staphylococcus pasteuri* SD)

| **Salt** | **Initial concentration (ppm)** | **Final concentration (ppm)** | **Percentage Accumulation** |
|---|---|---|---|
| **Nickel** | 90.57±1.15 | 65.68±4.11 | 27.48% |
| **Copper** | 88.44±3.16 | 62.48±3.85 | 29.35% |
| **Cadmium** | 84.46±4.03 | 70.37±2.73 | 19.84% |
| **Chromium** | 87.66±5.15 | 65.34±4.06 | 25.47% |
| **Lead** | 88.59±3.06 | 56.45±3.25 | 36.28% |
| **Iron** | 77.04±3.34 | 64.48±4.46 | 16.30% |

**Table 5.** Bioaccumulation by isolate B2 (*Sporosarcina saromensis* SAS)

| **Salt** | **Initial Concentration (ppm)** | **Final Concentration (ppm)** | **Percentage Accumulation** |
|---|---|---|---|
| **Nickel** | 88.82±3.16 | 59.35±3.68 | 33.18% |
| **Copper** | 81.63±4.04 | 66.13±5.15 | 18.98% |
| **Cadmium** | 78.29±3.64 | 63.75±3.98 | 18.57% |
| **Chromium** | 85.92±3.54 | 52.22±4.13 | 39.22% |
| **Lead** | 80.88±2.81 | 54.53±3.33 | 32.58% |
| **Iron** | 62.21±4.06 | 49.11±4.29 | 21.05% |

**Table 6.**
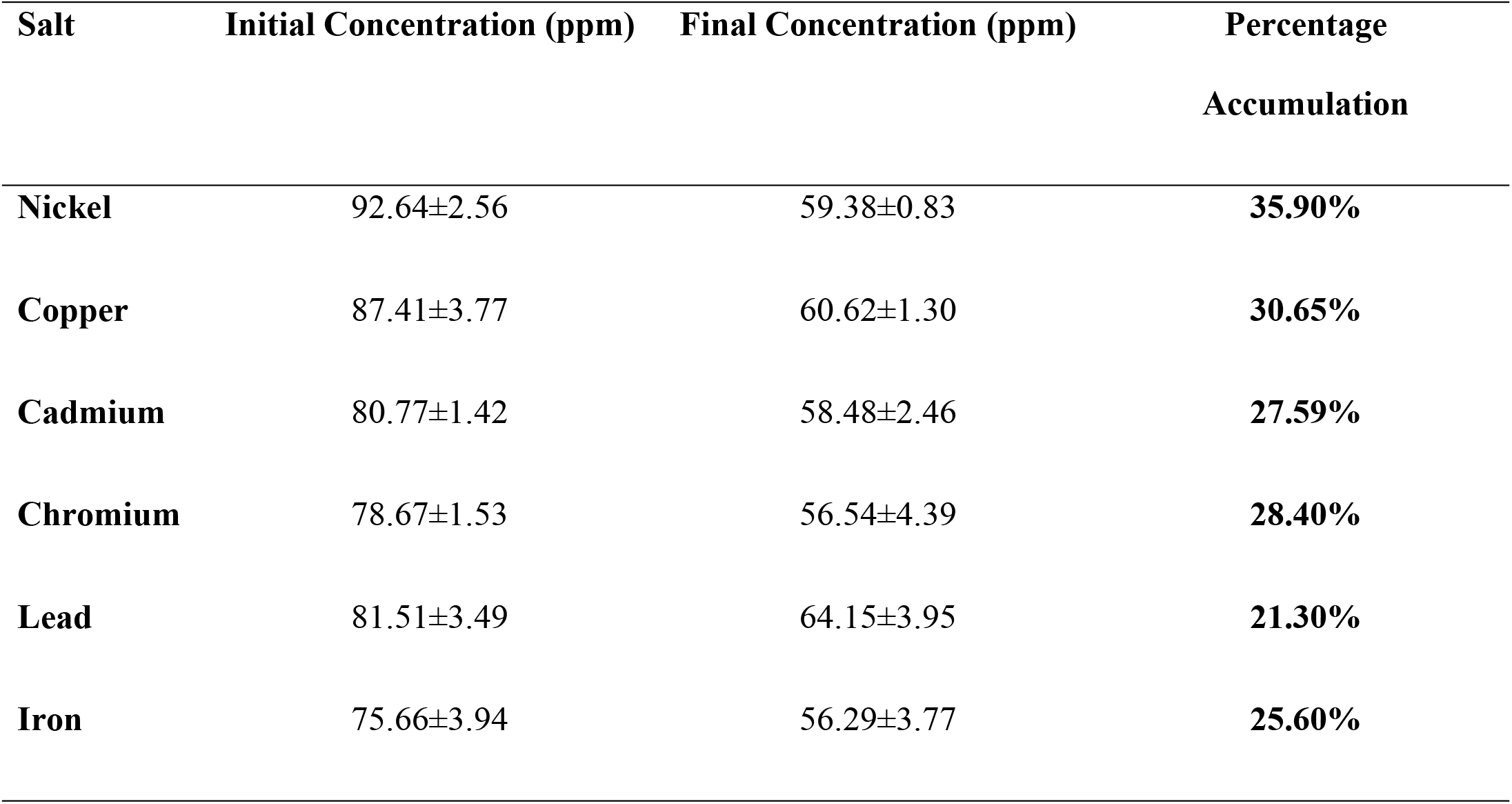
Bioaccumulation by isolate B3 (*Morganella morganii* SAS1)

### Identification of potential heavy metal remediating isolates

At the conclusion of this study, the three bacterial isolates, designated B1, B2, and B3, exhibiting heavy metal tolerance (HMT) and heavy metal remediation capacity were identified as *Staphylococcus pasteuri* SD, *Sporosarcina saromensis* SAS, and *Morganella morganii* SAS1, respectively by 16S rRNA gene sequencing. Their sequences were deposited in the GenBank database under accession numbers PZ182827.1, PZ179984.1, and PZ492667.1. BLASTN analysis revealed 99% sequence identity for all hits. Multiple sequence alignment was conducted using MEGA version 12.0 and demonstrated high similarity among the retrieved sequences from various bacterial species. A phylogenetic tree was subsequently constructed to delineate evolutionary relationships between the isolates and their closest known relatives (Figs 6, 7, and 8).

**Fig. 6:**
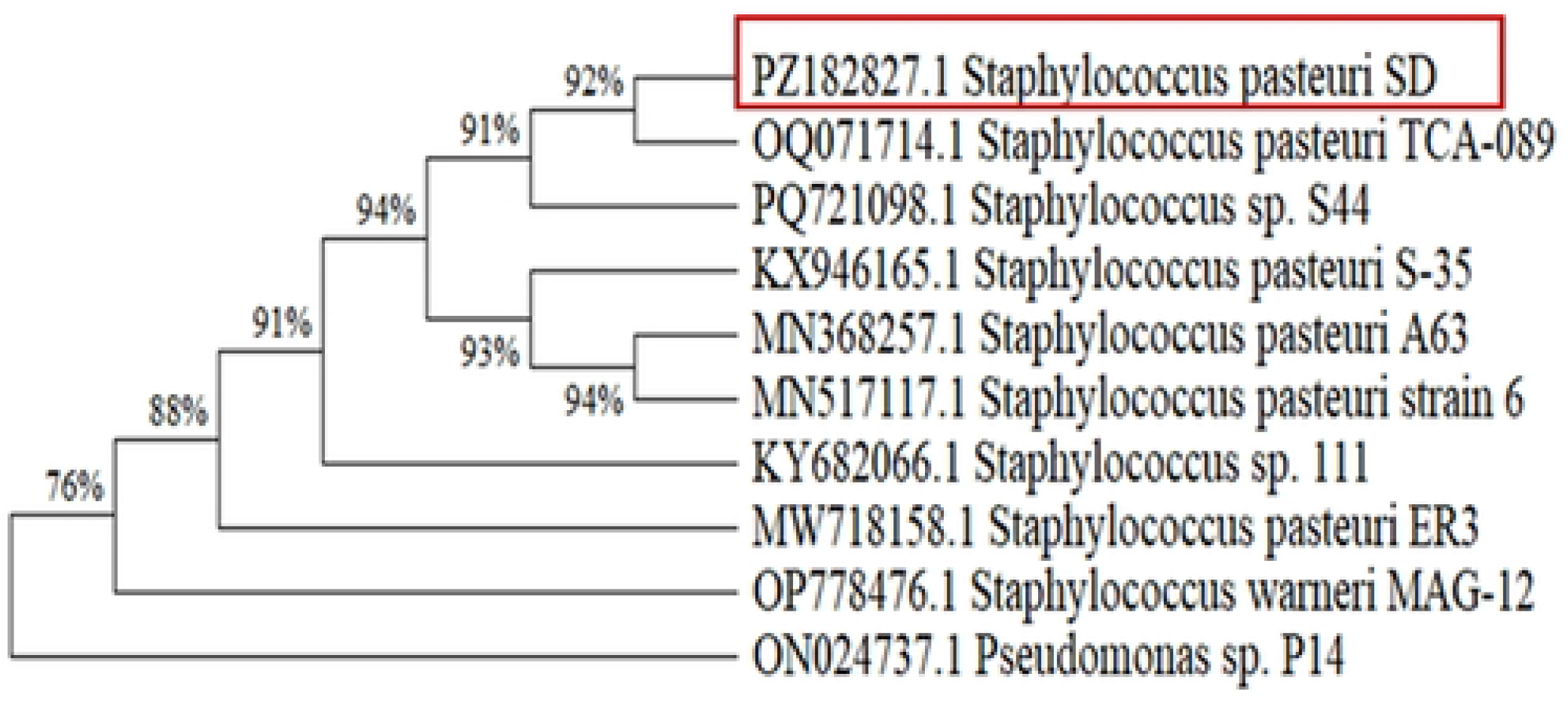
Phylogenetic tree of isolate BI *Staphylococcus pasteuri* SD. Partial 16S rRNA gene sequences from the HMR isolate *Staphylococcus pasteuri* SD and its closest related strains were aligned using ClustalW. A phylogenetic tree was then constructed using maximum-likelihood analysis with the Tamura-Nei model to assess their evolutionary relationships.

**Fig 7:**
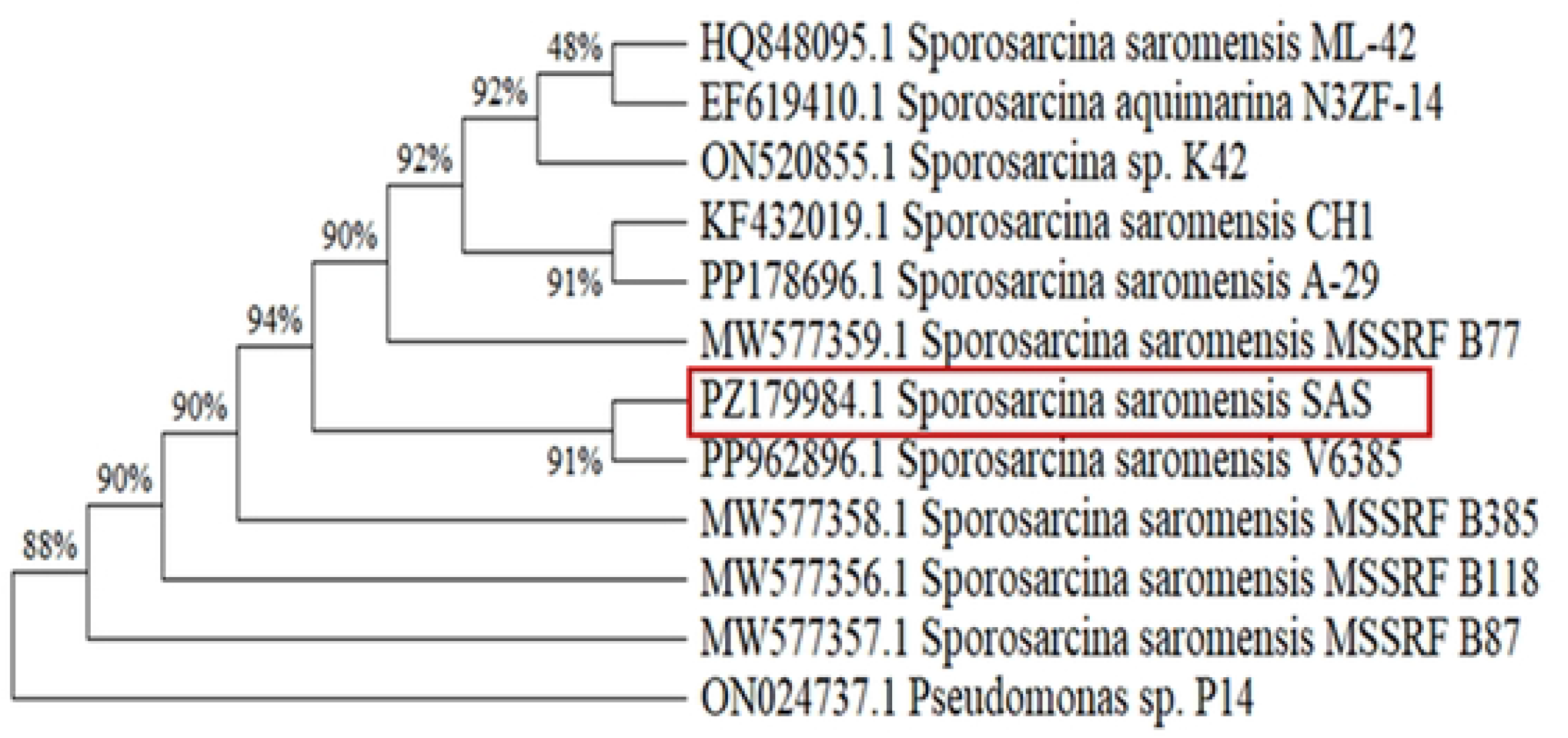
Phylogenetic tree of isolate B2 *Sporosarcina saromensis* SAS. Partial 16S rRNA gene sequences from the HMR isolate *Sporosarcina saromensis* SAS and its closest relatives were aligned using ClustalW. A phylogenetic tree was then constructed using maximum-likelihood analysis with the Tamura-Nei model to determine their evolutionary relationships.

**Fig 8:**
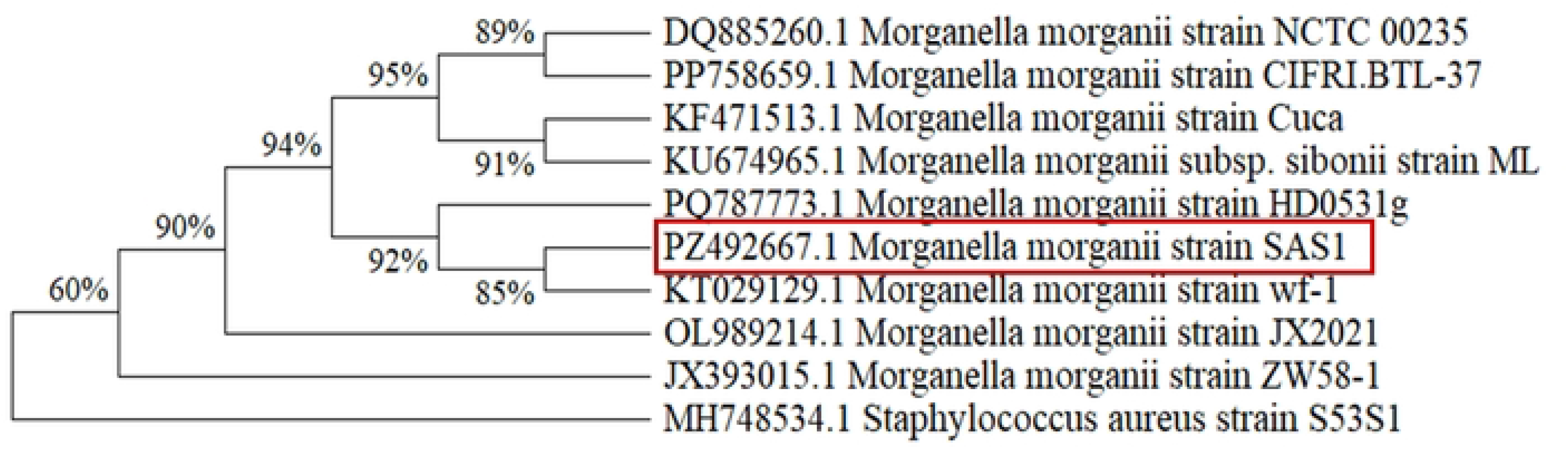
Phylogenetic tree of isolate B3 *Morganella morganii* SAS1. Partial 16S rRNA gene sequences of the HMR isolate *Morganella morganii* SAS1 and its nearest strains were aligned using ClustalW, and a phylogenetic tree was constructed through maximum-likelihood analysis employing the Tamura-Nei model to determine their evolutionary relationships.

Furthermore, the three isolates were characterized as *Staphylococcus pasteuri*, *Sporosarcina saromensis*, and *Morganella morganii* based on their morphological, physiological, cultural, and biochemical characteristics (S3 Table and S2, S3 and S4 Figs.) and by comparing these traits with the descriptions in the eighth edition of Bergey’s Manual of Determinative Bacteriology [48].

## Discussion

This study investigates the role of gut microbes in SMCF adaptability in the polluted Buriganga River and isolates bacteria from the SMCF’s gastrointestinal tract capable of remediating heavy metals. The Buriganga River is one of Bangladesh’s most contaminated waterways, and the rapid increase in SMCF populations raises significant ecological concerns in aquatic environments [5]. Studies show that to survive in heavily polluted waters, fish must adapt their physiology, potentially altering their gut microbial communities [8]. Prior research has examined SMCF adaptability in heavy-metal-contaminated environments, showing that substantial population growth is facilitated by the intestinal microbiota, particularly bacteria [8]. Despite the severe pollution, the dominance of Suckermouth catfish over edible fish species heightens ecological concerns. This study re-evaluated the pollution status of the Buriganga River at four sampling stations to confirm that collected fish samples originated from polluted sites. Water Quality Index (WQI) of each sampling location was assessed. The findings, detailed in Tables 1 and 2 and the S1 and S2 tables in the S1 file, showed elevated physicochemical parameters at all stations, with WQI values significantly exceeding the standard threshold of 100 (P < 0.05), indicating severe water quality deterioration. As previously reported, a WQI above 100 corresponds to grade E water, denoting very poor quality [45]. Such elevated indices suggest severely polluted conditions that likely disrupt normal aquatic habitats and threaten ecological stability. These results collectively indicate that the Buriganga River is heavily contaminated with various heavy metals from multiple point and non-point sources. Despite this, SMCF species are thriving, demonstrating remarkable adaptability to the harsh environment. The study also found heavy metal concentrations exceeding acceptable limits, with current levels comparable to or higher than those reported in earlier studies [49]. These high concentrations pose substantial risks to fish and other aquatic organisms. However, SMCFs demonstrated considerable resilience and proliferated despite heavy metal contamination, whereas most edible fish species declined sharply, with only a few remaining.

Regarding the microbial communities of the SMCF, the SMCF samples showed increased abundance of contaminant-remediating bacteria. Notably, *Bacillus, Paraclostridium, Klebsiella, Pseudomonas, Sporosarcina,* and *Morganella* were detected exclusively across all SMCF samples collected at the four Buriganga sites. The gut microbial community structure of the SMCFs from different locations in Buriganga was relatively stable, as shown in Fig1 and evidenced by the calculated Shannon diversity index of the SMCF gut microbiome. This stable gut microbiome profile of SMCFs, along with the minimal presence of other genera, indicates an associated microbial community. This result suggests that the dominant microbial community, along with other less abundant microbes in the gut of SMCF, may have contributed to the ability of SMCFs (*P. disjunctivus*) to thrive in contaminated river environments. Multiple studies have reported that *Bacillus, Paraclostridium, Staphylococcus Klebsiella, Pseudomonas, Sporosarcina, and Morganella* sp. have potential for heavy-metal bioremediation [41,50–57]. It is reported that the intestinal microbiota plays a crucial role in facilitating the survival and proliferation of SMCF in heavy-metal-contaminated aquatic environments [8]. This study also indicated that dominant taxa such as Actinobacteria, Firmicutes, and Proteobacteria can enzymatically transform heavy metals into less toxic forms, aiding SMCF survival [8]. These findings align with our study, suggesting that contaminant-tolerant microbes in SMCF reflect adaptive microbial colonization in response to environmental stress from heavy metals and other contaminants. Moreover, these findings suggest that the gut microbiota of SMCF holds considerable promise for heavy metal remediation and highlight the need for targeted bioremediation strategies. Additionally, we measured heavy metal levels in SMCF muscle tissue to evaluate their capacity for heavy metal accumulation. Alarmingly, we detected elevated levels of multiple heavy metals that exceeded recommended safety thresholds (Table 3). This finding also aligns with prior research showing that SMCF muscle tissue can accumulate heavy metals to critically elevated levels [58]. Together, these findings suggest that gut microbes in SMCFs may transform heavy metals into non-toxic forms and contribute to SMCFs’ remarkable adaptability in adverse aquatic environments.

Another objective of this study is to isolate and identify microbes with heavy-metal-remediating potential from the SMCF gut. Because SMCFs inhabit the heavily contaminated Buriganga River and harbor contaminant-remediating microbiota, their gut contents were targeted to isolate heavy metal-remediating bacteria. Three isolates (B1, B2, and B3) were selected for their potential to tolerate multiple heavy metals, as evidenced by their growth on LB medium supplemented with heavy metals. The maximum tolerance concentration (MTC) of multiple heavy metals by these isolates indicated that all three isolates exhibited distinct tolerance responses (Fig2). The isolates demonstrated a wide range of tolerance to multiple heavy metals, with the highest tolerable concentration recorded at 1000 ppm for at least one metal (Fig2). Previous research reported that bacteria tolerant of one metal often tolerate several others due to shared toxic mechanisms among heavy metals [41]. A group of scientists isolated a *Bacillus* sp. from heavy metal-contaminated soil that exhibited resistance to a broad spectrum of heavy metals [41]. In the present study, isolates B1, B2, and B3 demonstrated resistance to multiple heavy metals, suggesting their capacity to tolerate various heavy-metal stresses and to accumulate and/or remediate different heavy metals. We also examined the growth patterns of isolates B1, B2, and B3 under heavy-metal stress conditions. All isolates showed robust growth under multiple heavy-metal stress conditions, comparable to or even exceeding growth in the control condition (lacking metal supplementation) (Fig 3, 4, and 5). Prior research indicated that *Cellulosimicrobium* sp. exhibited variable growth over 48 hours in metal-stressed environments, demonstrating its potential for multiple heavy-metal tolerance [39]. Another study identified heavy-metal tolerance traits, including resistance to Cd²⁺, Cr⁶⁺, Ni²⁺, and Co²⁺, based on bacterial growth under heavy-metal–supplemented conditions [41]. These findings are consistent with our current results. Therefore, the selected isolates can tolerate varying concentrations of multiple heavy metals and have remediation potential. Several studies have noted that organisms resistant to multiple heavy metals could serve as promising candidates for bioremediation applications [39,59]. To validate this potential, we conducted a dedicated bioremediation assay, which showed that all three isolates exhibited varying percentages of bioremediation efficacy against different metals (Tables 4, 5, and 6). Among the tested isolates, B1 showed the highest lead accumulation (36.28%), while B2 showed the highest chromium accumulation (39.22%). In parallel, B3 displayed the highest nickel accumulation (35.90%). These findings suggest that the microbes isolated in this study are promising candidates for remediating heavy metal-contaminated sites. Additionally, these results imply that a microbial consortium combining these isolates could be developed to enhance bioremediation efficiency at contaminated sites.

Finally, we identified all three isolates through molecular characterization, including 16S rRNA gene sequencing, BLASTN analysis, and phylogenetic tree construction (Figures 6, 7 & 8). We also assessed their cultural, physiological, morphological, and biochemical characteristics using complementary approaches (S3 Table and S2, S3, and S4 Figs). The isolates B1, B2, and B3 were characterized as *Staphylococcus pasteuri* SD (Accession Number: PZ182827.1), *Sporosarcina saromensis* SAS (Accession Number: PZ179984.1), and *Morganella morganii* SAS1 (Accession Number: PZ492667), respectively. Next-generation sequencing (NGS) analysis of the SMCF gut microbiota further confirmed that these bacteria were stably present in the gut. Previous research indicates that bacterial strains such as *Bacillus cereus*, *Staphylococcus aureus*, *Escherichia coli*, and *Proteus* spp., isolated from a Nigerian cement industry site, display notable resistance to heavy metal exposure, suggesting that these microorganisms possess adaptive traits suitable for polluted environments, thereby making them promising candidates for bioremediation [52]. Moreover, other studies have shown that *Staphylococcus* species resist multiple heavy metals, underscoring their potential in bioremediation initiatives [53]. Chinese researchers have investigated the bioremediation capacity of *Sporosarcina saromensis* M52, reporting its ability to tolerate and detoxify chromium [51]. Additionally, previous studies have shown that *Morganella morganii*, which tolerates nickel, has significant potential to enhance plant growth and photosynthetic efficiency in nickel-contaminated soils[57]. The findings of the present study corroborate earlier research, as the isolates—*Staphylococcus pasteuri* SD, *Sporosarcina saromensis* SAS, and *Morganella morganii* SAS1—also demonstrate tolerance to various heavy metals and possess effective bioremediation capabilities. These results underscore that genera isolated from the SMCF gut harbor multiple traits conferring heavy-metal resistance, highlighting their potential utility in bioremediation applications.

## Conclusion

In summary, this study demonstrated that the gut microbiome of SMCFs collected from the heavily polluted Buriganga River in Bangladesh exhibits a high prevalence of bacteria capable of remediating various contaminants, including heavy metals. These characteristics may facilitate contaminant management and thereby enhance their notable adaptability in such polluted settings. This finding is further supported by bacteria isolated from the gut of SMCFs, which demonstrate an extraordinary capacity to remediate multiple heavy metals. Three identified strains—*Staphylococcus pasteuri* SD, *Sporosarcina saromensis* SAS, and *Morganella morganii* SAS1— show significant potential for heavy-metal bioremediation. Their high tolerance and resilience to heavy-metal stress make them promising candidates for bioremediation of heavy-metal-contaminated sites and suggest developing microbial consortia that use these isolated bacteria to remediate not only heavy metals but also other contaminants.

## Acknowledgments

We gratefully acknowledge support from the Department of Microbiology, University of Chittagong, Bangladesh, and the Research and Publication Cell, University of Chittagong, Bangladesh, in conducting the research.

## Author contributions

**Conceptualization:** Sujan Dey

**Data curation:** Sumaita Alam Sikder, Sujan Dey

**Formal analysis:** Sujan Dey, Sumaita Alam Sikder

**Investigation:** Sumaita Alam Sikder, Mohsina Rahman Epa, Sujan Kanti Das

**Methodology:** Sumaita Alam Sikder, Sujan Dey

**Funding acquisition:** Sujan Dey, Md. Zobaidul Alam

**Project administration:** Sujan Dey, Md. Zobaidul Alam

**Resources:** Sujan Dey

**Supervision:** Sujan Dey

**Validation:** Sujan Dey. Md. Zobaidul Alam

**Writing – original draft:** Sujan Dey, Sumaita Alam Sikder

**Writing – review & editing:** Sujan Dey, Sumaita Alam Sikder, Md. Zobaidul Alam

## Supporting information

**S1 Table: Physicochemical parameters of water samples.**

**S2 Table: Concentration of different anions in water**

**S3 Table: Cultural, physiological, morphological, and biochemical characteristics of isolates.** The cultural, physiological, morphological, and biochemical characteristics were compared with the descriptions in the eighth edition of Bergey’s Manual of Determinative Bacteriology to confirm isolate B1 as *Staphylococcus pasteuri*, B2 as *Sporosarcina saromensis*, and B3 as *Morganella morganii*

**S1 Fig: Representative SMCF (*Pterigoplicthys disjunctivus*) samples collected from Buriganga River.**

**S2 Fig: Colony of isolate B1 (*Staphylococcus pasteuri* SD) grown in LB agar supplemented with metal salt.**

**S3 Fig: Colony of isolate B2 (*Sporosarcina saromensis* SAS) grown in LB agar supplemented with metal salt.**

**S4 Fig: Colony of isolate B3 (*Morganella morganii* SAS1) grown in LB agar supplemented with metal salt.**

## References

1. Nico L, Butt P, Johnston G, Jelks H, Kail M, Walsh S. Discovery of South American suckermouth armored catfishes (Loricariidae, Pterygoplichthys spp.) in the Santa Fe River drainage, Suwannee River basin, USA. BioInvasions Rec. 2012;1(3):179–200. doi:10.3391/bir.2012.1.3.04

2. Wu GD, Chen J, Hoffmann C, Bittinger K, Chen YY, Keilbaugh SA, et al. Linking Long-Term Dietary Patterns with Gut Microbial Enterotypes. Science. 2011;334(6052):105–8. doi:10.1126/science.1208344

3. Mack RN, Simberloff D, Mark Lonsdale W, Evans H, Clout M, Bazzaz FA. BIOTIC INVASIONS: CAUSES, EPIDEMIOLOGY, GLOBAL CONSEQUENCES, AND CONTROL. Ecol Appl. 2000;10(3):689–710. doi:10.1890/1051-0761(2000)010[0689:bicegc]2.0.co;2

4. Hoover JJ, Killgore KJ, Cofrancesco Jr AF. Suckermouth catfishes: threats to aquatic ecosystems of the United States? 2004; : https://tsusinvasives.org/dotAsset/a6574267-0f10-4d47-844e-1065e2d50a20.pdf

5. Department of Fisheries Resource Management, Faculty of Fisheries, Chattogram Veterinary and Animal Sciences University, Chattogram, Bangladesh, Sarkar A, Rana S, Bhowmik P, Hasan MdN, Shimul SA, et al. A Review of Suckermouth Armoured Catfish (Siluriformes: Loricariidae) Invasion, Impacts and Management: Is Its Invasion a Threat to Bangladesh’s Fisheries Sector? Asian Fish Sci. 2023;36(3). doi:10.33997/j.afs.2023.36.3.002

6. Rana MM, Mahmud S, Siam KHH, Chowdhury HA, Das SR, Islam M, et al. Invasive Suckermouth catfishes (Siluriformes: Loricariidae) in Bangladesh wetlands: A review on the present status and challenges. Bangladesh J Zool. 2023;51(3):371–96.

7. Karunarathna D, Amarasinghe AT, Ekanayake E. Observed predation on a suckermouth catfish (Hypostomus plecostomus) by a water monitor (Varanus salvator) in Bellanwila-Attidiya Sanctuary. Biawak. 2008;2(1):37–9.

8. Elfidasari D, Rijal MS, Shalsabilla SE, Rahma Fadila DS, Cici A, Pikoli MR, et al. Intestinal bacteria diversity of suckermouth catfish (*Pterygoplichthys pardalis*) in the Cd, Hg, and Pb contaminated Ciliwung River, Indonesia. Heliyon. 2023;9(4):e14842. doi:10.1016/j.heliyon.2023.e14842

9. Hossain I, Mamun ASA, Bharati P, Hossain G. Prevalence and risk factors of malnutrition among primary school children from 1998-2017: A systematic review and meta-analysis. 2018.

10. Mori S, Nakamura T. An evolutionarily conserved odontode gene regulatory network underlies head armor formation in suckermouth armored catfish. bioRxiv; 2021 doi:10.1101/2021.06.21.449322

11. Kamal MM, Malmgren-Hansen A, Badruzzaman ABM. Assessment of pollution of the River Buriganga, Bangladesh, using a water quality model. Water Sci Technol. 1999; Water Quality and its Management 40(2):129–36. doi:10.1016/S0273-1223(99)00474-6

12. Pasha A, Abdillahi MM, Rahman SM, Mozumder S, Chowdhury AH, Fuente JAD, et al. STUDIES ON PHYSICOCHEMICAL PROPERTIES OF BURIGANGA RIVER WATER AND THE VEGETATION COVERAGE OF SURROUNDING AREA, DHAKA, BANGLADESH. Sci IntLahore. 2022;34(2):73–8.

13. Kibria MG, Kadir MN, Alam S. Buriganga river pollution: its causes and impacts. In: International Conference on Recent Innovation in Civil Engineering for Sustainable Development (IICSD-2015). 2015. p. 323–8.

14. Tilzer MM, Khondker M. Hypertrophic and polluted freshwater ecosystems: Ecological basis for water resource management. Dept Bot Dhaka Univ Bangladesh. 1993.

15. Bhuiyan MAH, Dampare SB, Islam MA, Suzuki S. Source apportionment and pollution evaluation of heavy metals in water and sediments of Buriganga River, Bangladesh, using multivariate analysis and pollution evaluation indices. Environ Monit Assess. 2015;187(1). doi:10.1007/s10661-014-4075-0

16. Ahmed K, Ahamed S, Rahman S, Haque R, Islam M. Heavy Metals Concentration in Water, Sediments and their Bioaccumulations in Some Freshwater Fishes and Mussel in Dhaleshwari River, Bangladesh.

17. Reza A, Yousuf TB. Impacts of Waste Dumping on Water Quality in the Buriganga River, Bangladesh and Possible Mitigation Measures.

18. Hemmat-Jou MH, Safari-Sinegani AA, Mirzaie-Asl A, Tahmourespour A. Analysis of microbial communities in heavy metals-contaminated soils using the metagenomic approach. Ecotoxicology. 2018;27(9):1281–91. doi:10.1007/s10646-018-1981-x

19. Wang J, Chen C. Biosorption of heavy metals by *Saccharomyces cerevisiae*: A review. Biotechnol Adv. 2006 Sep 1;24(5):427–51. doi:10.1016/j.biotechadv.2006.03.001

20. Vijayaraj AS, Mohandass C, Joshi D. Microremediation of tannery wastewater by siderophore producing marine bacteria. Environ Technol. 2020;41(27):3619–32. doi:10.1080/09593330.2019.1615995 PubMed PMID: 31070993.

21. Sessitsch A, Kuffner M, Kidd P, Vangronsveld J, Wenzel WW, Fallmann K, et al. The role of plant-associated bacteria in the mobilization and phytoextraction of trace elements in contaminated soils. Soil Biol Biochem. 2013;60:182–94.

22. Teles YV, De Castro LM, Sargentini Junior É, Do Nascimento AP, Da Silva HA, Costa RS, et al. Potential of Bacterial Isolates from a Stream in Manaus-Amazon to Bioremediate Chromium-Contaminated Environments. Water Air Soil Pollut. 2018;229(8). doi:10.1007/s11270-018-3903-1

23. Ahemad M, Kibret M. Recent Trends in Microbial Biosorption of Heavy Metals: A Review. Biochem Mol Biol. 2013;1(1):19. doi:10.12966/bmb.06.02.2013

24. Abdu N, Abdullahi AA, Abdulkadir A. Heavy metals and soil microbes. Environ Chem Lett. 2017;15(1):65–84. doi:10.1007/s10311-016-0587-x

25. Alsafran M, Saleem MH, Al Jabri H, Rizwan M, Usman K. Principles and Applicability of Integrated Remediation Strategies for Heavy Metal Removal/Recovery from Contaminated Environments. J Plant Growth Regul. 2023;42(6):3419–40.doi:10.1007/s00344-022-10803-1

26. Verma S, Kuila A. Bioremediation of heavy metals by microbial process. Environ Technol Innov. 2019;14:100369. doi:10.1016/j.eti.2019.100369

27. Lun H, Yang W, Zhao S, Jiang M, Xu M, Liu F, et al. Altered gut microbiota and microbial biomarkers associated with chronic kidney disease. MicrobiologyOpen. 2019;8(4):e00678. doi:10.1002/mbo3.678

28. Sultana S, Khan MdN, Hossain MS, Dai J, Rahman MS, Salimullah Md. Community Structure and Functional Annotations of the Skin Microbiome in Healthy and Diseased Catfish, Heteropneustes fossilis. Front Microbiol. 2022;13:856014. doi:10.3389/fmicb.2022.856014

29. Quast C, Pruesse E, Yilmaz P, Gerken J, Schweer T, Yarza P, et al. The SILVA ribosomal RNA gene database project: improved data processing and web-based tools. Nucleic Acids Res. 2013;41(D1):D590–6. doi:10.1093/nar/gks1219

30. Dey S, Awata T, Mitsushita J, Zhang D, Kasai T, Matsuura N, et al. Promotion of biological nitrogen fixation activity of an anaerobic consortium using humin as an extracellular electron mediator. Sci Rep. 2021;11(1):6567. doi:10.1038/s41598-021-85955-3

31. Akan JC. Physical and Chemical Parameters in Abattoir Wastewater Sample, Maiduguri Metropolis, Nigeria.

32. Carpenter JH. The Accuracy of the Winkler Method for Dissolved Oxygen Analysis. Limnol Oceanogr. 1965;10(1):135–40. doi:10.4319/lo.1965.10.1.0135

33. Boyles W. Technical Information Series, Booklet No. 9.

34. Dey S, Das J, Manchur MA. Studies on Heavy Metal Pollution of Karnaphuli River, Chittagong, Bangladesh.

35. Tahmina B, Sujan D, Karabi R, Hena MKA, Amin KR, Sharmin S. Assessment of Surface Water Quality of the Turag River in Bangladesh. Res J Chem Environ. 2018.

36. Kassim NSA, A. I. S. M. Ghazali S, Liyana Bohari F, A. Z. Abidin N. Assessment of heavy metals in wastewater plant effluent and lake water by using atomic absorption spectrophotometry. Mater Today Proc. 2022 Jan 1;8th International Conference on Solid State Science & Technology 66:3961–4. doi:10.1016/j.matpr.2022.04.671

37. Leblanc PJ, Jackson AL. Dry Ashing Technique for the Determination of Arsenic in Marine Fish. J Assoc Off Anal Chem. 1973;56(2):383–6. doi:10.1093/jaoac/56.2.383

38. Niazi SB, Littlejohn D, Halls DJ. Rapid partial digestion of biological tissues with nitric acid for the determination of trace elements by atomic spectrometry. Analyst. 1993;118(7):821–5. doi:10.1039/AN9931800821

39. Bhati T, Gupta R, Yadav N, Singh R, Fuloria A, Waziri A, et al. Assessment of Bioremediation Potential of Cellulosimicrobium sp. for Treatment of Multiple Heavy Metals. Microbiol Biotechnol Lett. 2019;47(2):269–77. doi:10.4014/mbl.1808.08006

40. Kalaimurugan D, Balamuralikrishnan B, Durairaj K, Vasudhevan P, Shivakumar MS, Kaul T, et al. Isolation and characterization of heavy-metal-resistant bacteria and their applications in environmental bioremediation. Int J Environ Sci Technol. 2020;17(3):1455–62. doi:10.1007/s13762-019-02563-5

41. Samanta A, Bera P, Khatun M, Sinha C, Pal P, Lalee A, et al. An investigation on heavy metal tolerance and antibiotic resistance properties of bacterial strain Bacillus sp. isolated from municipal waste. 2012.

42. Acinas SG, Klepac-Ceraj V, Hunt DE, Pharino C, Ceraj I, Distel DL, et al. Fine-scale phylogenetic architecture of a complex bacterial community. Nature. 2004;430(6999):551–4. doi:10.1038/nature02649

43. Kumar S, Stecher G, Li M, Knyaz C, Tamura K. MEGA X: Molecular Evolutionary Genetics Analysis across Computing Platforms. Battistuzzi FU, editor. Mol Biol Evol. 2018;35(6):1547–9. doi:10.1093/molbev/msy096

44. Saitou N, Nei M. The neighbor-joining method: a new method for reconstructing phylogenetic trees. Mol Biol Evol. 1987;4(4):406–25. doi:10.1093/oxfordjournals.molbev.a040454

45. Tyagi S, Sharma B, Singh P, Dobhal R. Water Quality Assessment in Terms of Water Quality Index. Am J Water Resour. 2020;1(3):34–8. doi:10.12691/ajwr-1-3-3

46. World Health Organization (2008). Guidelines for drinking-water quality [electronic resource]: incorporating 1st and 2nd addenda,Vol.1, recommendations, 3rd ed. World Health Organization. https://iris.who.int/handle/10665/204411

47. FAO, 1983. (Food and Agriculture Organization), Compilation of legal limits for hazardous substances in fish and fishery products, FAO Fishery Circular No. 464, pp. 5–100.

48. Balows A. Bergey’s Manual of Determinative Bacteriology. Eighth Edition. Am J Public Health. 1975;65(3):315–315. doi:10.2105/AJPH.65.3.315-a

49. Hoque M, Deb P. Assessment of Physicochemical Water Quality Parameters and Heavy Metals Concentration in Water Samples from Buriganga River Adjacent to Dhaka City. J Environ Sci Nat Resour. 2017;9(2):97–104. doi:10.3329/jesnr.v9i2.32164

50. M B B, Shini S, Mohan M. Mercury Biosorption and merA Gene Identification in an Anaerobic Bacterium Paraclostridium bifermentans B1 Isolated from Cochin Estuary. 2019.

51. Ran Z, Bi W, Tao CQ, Xia LX, Min L, Dong H, et al. Bioremediation of Hexavalent Chromium Pollution by Sporosarcina saromensis M52 Isolated from Offshore Sediments in Xiamen, China.

52. Martins AL, Silas TV, Abah MA, Adebisi AK, Sunday AM, Emochonne RY, et al. Isolation, identification, and characterization of heavy metal-resistant bacteria from soil samples collected at a cement company in Nigeria. Asian J Trop Biotechnol. 2024;21(1). doi:10.13057/biotek/c210103

53. Suresh G, Balasubramanian B, Ravichandran N, Ramesh B, Kamyab H, Velmurugan P, et al. Bioremediation of hexavalent chromium-contaminated wastewater by Bacillus thuringiensis and Staphylococcus capitis isolated from tannery sediment. Biomass Convers Biorefinery. 2021;11(2):383–91. doi:10.1007/s13399-020-01259-y

54. Atuchin VV, Asyakina LK, Serazetdinova YR, Frolova AS, Velichkovich NS, Prosekov AY. Microorganisms for Bioremediation of Soils Contaminated with Heavy Metals. Microorganisms. 2023;11(4):4. doi:10.3390/microorganisms11040864

55. Rajbanshi A. Study on Heavy Metal Resistant Bacteria in Guheswori Sewage Treatment Plant. Our Nat. 2008;6(1):1. doi:10.3126/on.v6i1.1655

56. Abbas S, Zulfiqar S, Arshad M, Khalid N, Hussain A, Ahmed I. Molecular characterization of heavy metal-tolerant bacteria and their potential for bioremediation and plant growth promotion. Front Microbiol. 2025;16. doi:10.3389/fmicb.2025.1644466

57. Naqqash T, Aziz A, Baber M, Shahid M, Sajid M, Emanuele R, et al. Metal-tolerant morganella morganii isolates can potentially mediate nickel stress tolerance in Arabidopsis by upregulating antioxidative enzyme activities. Plant Signal Behav. 2024;19(1):2318513. doi:10.1080/15592324.2024.2318513 PubMed PMID: 38526224.

58. Awal MdR, Moniruzzaman Md, Hossen MdN, Amin A, Said MdA, Islam MdS, et al. Nutrient profile and heavy metals content of an exotic fish Hypostomus plecostomus in Bangladesh: Health risk assessment. Aquat Invertebr Ecosyst Res. 2024;01(01):14–20. doi:10.69517/aier.2024.01.01.0003

59. Dinku A, Jiru TM. Isolation, screening, and characterization of heavy metal-resistant bacteria from solid waste dumping sites in Central Gondar, Northwest, Ethiopia. Sci Prog. 2025;108(1). doi:10.1177/00368504251315807

